# Visual cortex anodal tDCS does not alter reading performance for Chinese presented character-by-character to normal peripheral vision

**DOI:** 10.1101/2023.11.19.567752

**Authors:** Anqi Lyu, Andrew E Silva, Benjamin Thompson, Larry Abel, Allen MY Cheong

## Abstract

Visual cortex transcranial direct current stimulation (tDCS) reduces crowding in normal peripheral vision and may improve reading of English words in patients with macular degeneration. Given the different visual requirements of reading English words and Chinese characters, the effect of tDCS on peripheral reading performance in English might be different from Chinese. This study recruited seventeen participants (59 to 73 years of age) with normal vision and tested the hypothesis that anodal tDCS would improve reading of Chinese characters presented at 10° eccentricity compared with sham stimulation. Chinese sentences of different print sizes and exposure durations were presented one character at a time 10° below or to the left of fixation, and the individual critical print size (CPS) - the smallest print size eliciting the maximum reading speed (MRS) was determined. Reading accuracies for characters presented 0.2 logMAR smaller than the CPS were measured before, during, 5 mins, and 30 mins after receiving active or sham anodal visual cortex tDCS. Participants completed both the active and sham sessions in a random order following a double-blind, within-subject design. No effect of active anodal-tDCS on reading accuracy was observed, implying that a single session of tDCS did not improve Chinese character reading in normal peripheral vision. This may suggest that tDCS does not significantly reduce the crowding elicited within a single Chinese character. However, the effect of tDCS on between-character crowding is yet to be determined.

## Introduction

Reading speed is faster when using central vision compared with peripheral vision^1-3^, possibly due to reduced spatial resolution^4^, poorer eye movement control^5,6^, restricted visual span^7^, and susceptibility to crowding^1,8^. Though few studies have investigated the factors affecting Chinese reading performance using peripheral vision, our recent findings have revealed systematic impairments in peripheral Chinese reading, including slower temporal processing speeds and reduced spatial visual spans^9^. Understanding the peripheral reading performance in readers with normal vision may help generate better rehabilitation solutions for the low-vision community.

Clinically, reading difficulty is a major concern for patients with age-related macular degeneration (AMD), who are forced to rely on para-central or peripheral vision because of the presence of a central scotoma. Studies of interventions designed to improve normal peripheral vision not only enhance our understanding of the effects of interventions on reading performance but may also facilitate their future adoption for patients with AMD. For example, increasing English letter spacing (within a particular range) to reduce crowding improved peripheral reading speed^1,10^. Multiple training sessions on a crowded letter identification task also improved peripheral English reading performance^11^.

Recently, non-invasive brain stimulation (NIBS), which enables the modulation of neural activity in targeted superficial areas of the human brain, is emerging as a promising tool for vision enhancement and rehabilitation. Anodal transcranial direct current stimulation (a-tDCS), a common NIBS technique^12^, has been demonstrated to improve a range of visual functions. Results from our recent meta-analysis illustrated that visual cortex a-tDCS improved contrast sensitivity, visual evoked potential amplitude (an index of cortical excitability), and crowding in peripheral vision among normally sighted individuals^13^. TDCS involves a weak 1 - 2 mA electrical current delivered through two head-mounted electrodes (the anode and cathode) and induces regional changes in cortical excitability and neurotransmitter concentrations that outlast the duration of stimulation (see Roche et al.^14^ for a review). Several studies have explored the effect of visual cortex a-tDCS on normal peripheral vision. Reinhart et al. showed that a single 20-min session of a-tDCS applied to the visual cortex significantly improved vernier acuity at 5° eccentricity in 20 normally-sighted subjects^15^. Furthermore, the acute effect of a-tDCS on collinear lateral inhibition, which is one of the low-level inhibitory mechanisms contributing to visual crowding, was examined in 13 subjects with normal vision. The participants detected a central target presented 6° left of fixation which was vertically flanked by 2 colinear Gabor patches. Lateral inhibition was significantly reduced following 20 mins of visual cortex a-tDCS whereas sham stimulation had no effect. Raveendran et al. proposed that the early stages of visual processing of peripheral stimuli could be enhanced by a-tDCS, potentially benefiting the high-level visual processing of crowded stimuli^16^. Indeed, Chen et al. demonstrated that the contrast threshold for recognizing an English letter at 10° eccentricity crowded by 2 random letters decreased (an improvement of 23%) after 20 mins of active a-tDCS, but not sham tDCS^17^.

The reduction of crowding in peripheral vision resulting from visual cortex a-tDCS suggests that this intervention may improve peripheral reading. In the first study to address this question, Silva et al. conducted a randomized, within-subjects experiment to examine the acute effects of visual cortex a-tDCS on English and Chinese reading performance in patients with binocular central vision loss. They found that 20 mins of a-tDCS tended to improve reading accuracy for English compared with sham stimulation. However, reading accuracy for Chinese did not change with a-tDCS^18^. The authors concluded that the effects of visual cortex a-tDCS on peripheral reading varied for different writing systems. However, reading abilities vary widely in AMD patients depending on their remaining vision and the location of their preferred retinal locus^19-21^. It has also been argued that AMD is accompanied by negative peripheral retinal manifestations because of the increased prevalence of peripheral autofluorescence abnormalities (see Pivovar & Oellers^22^ for a recent review). Hence, the remaining peripheral visual function might vary substantially across patients. Both of these factors might mask an effect of a-tDCS on reading in patients with AMD. Furthermore, in Silva et al.’s study, English reading performance was evaluated through a word-by-word presentation of sentences, whereas Chinese reading was examined by presenting sentences using a character-by-character approach^18^. Chinese, as an orthographic language, is characterized by logographic characters exhibiting varying spatial complexities. Chinese reading involves both internal (within-character) and external (between-character) crowding. It remains unclear whether the different effect of a-tDCS attributes to the language or methodological differences in reading measures. Therefore, the current study examined the effect of visual cortex a-tDCS on Chinese reading performance at a fixed peripheral location in a group of older adults with normal vision, using the same method described in Silva et al^18^. We assessed reading performance at 10° below and 10° left of fixation before and after applying a-tDCS. In contrast to the null results found in AMD patients, we hypothesized that Chinese peripheral reading performance would be improved after active visual cortex a-tDCS relative to the sham a-tDCS in older adults with normal vision.

## Methods

### Subjects

17 healthy older subjects (aged between 59 and 73 years) with normal or corrected-to-normal vision were recruited from community centers and social media. All subjects spoke and read traditional Chinese as their first language. Eligibility criteria included no history of any ophthalmic, neurological, or psychiatric illness, no taking medications that could affect reading performance, or any contraindication to brain stimulation such as epileptic seizure or metal/electronic implants in the head. Participants were divided into 2 groups: 1) reading at 10° left visual field (n=10), and 2) reading at 10° inferior visual field (n=7). The rationale for adopting two peripheral locations was because these are the two most frequent para-central locations adopted by AMD patients fixating at a central cross (33.7% and 39% respectively)^23^. Demographic data are summarized in Table 1. This experiment was approved by The Department Research Committee of the School of Optometry of The Hong Kong Polytechnic University (HSEARS20190716005). All subjects gave written informed consent. The study followed the tenets of the Declaration of Helsinki.

**Table 1.** Demographic data of the subjects. LogMAR = logarithm of the minimum angle of resolution measured using the Early Treatment Diabetic Retinopathy Study (ETDRS) chart at 4m (distance) and 40cm (near). Two-tailed student’s t-test or Mann-Whitney U test was used to compare the difference between groups. Mean ± standard deviation (SD) was calculated across groups.

|  | 10° left visual field<br>(n=10) | 10° inferior visual field<br>(n=7) | p-value |
| --- | --- | --- | --- |
| Age (years) | 65.73 ± 4.69<br>(59 – 73) | 64.68 ± 4.05<br>(60 – 71) | t(14)=0.49; p=0.63 |
| Gender | F: 80%<br>M: 20% | F: 43%<br>M: 57% | U=22; p=0.16 |
| Best corrected visual acuity (logMAR) | Distance: 0.06 ± 0.06<br>Near: 0.13 ± 0.09 | Distance: -0.01 ± 0.07<br>Near: 0.06 ± 0.07 | t(13)=2.00; p=0.07<br>U=17; p=0.08 |
| RSVP reading speed<br>(number of characters per min) | 159.30 ± 54.90 | 146.59 ± 59.09 | t(12)=0.45; p=0.66 |

### Experimental design

A double-blind, cross-over design was employed. Subjects were invited to The Hong Kong Polytechnic University Optometry Research Clinic and completed evaluations on 3 occasions. All tests were performed using their right eye with full refractive correction and appropriate near addition while the left eye was occluded with an eye patch.

Reading performance was assessed using rapid serial visual presentation (RSVP) which involved character-by-character presentation of sentences on a screen^1,7,24,25^. We adopted the same approach detailed in Silva et al.^18^. Chinese characters were generated using PsychoPy 2020.1.3^26^ and presented on a 24-inch LCD monitor (BENQ xl2540, 120 Hz refresh rate, 1920 x 1080 resolution). Subjects were seated 65 cm in front of the monitor with a chinrest and forehead bar stabilizing their head position. Subjects were asked to maintain fixation on a central cross. First, a short mask of “XXX” was presented at either 10° left of fixation or below fixation. Then, a randomly selected sentence replaced the mask and was presented one character at a time at the same location and with the same size as the initial mask (Fig.1). The sentence was selected randomly from a pool of 605 Chinese sentences comprising 15 characters. Subjects verbally reported all recognized characters. Eye movements were monitored using an infrared video eye-tracking system (Eyelink Portable Duo, SR Research, Scarborough, ONT, Canada). Trials were replaced if fixation deviated by more than 1°.

**Fig. 1.**
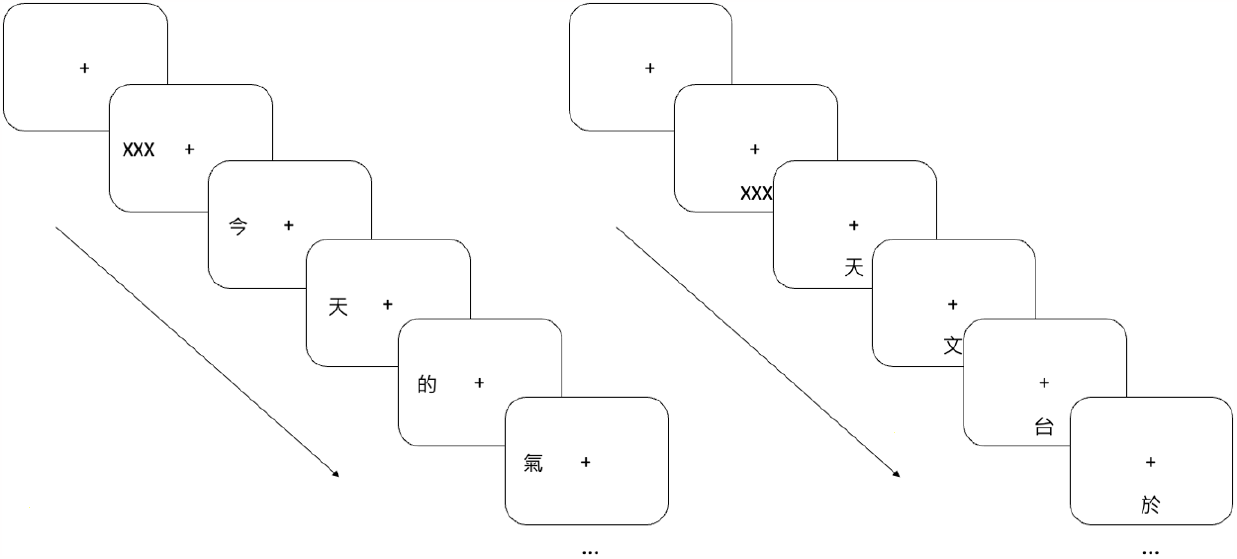
Rapid Serial Visual Presentation (RSVP). Character-by-character sentences were presented 10° left (left panel) or 10° below (right panel) the fixation cross. Subjects verbally reported the characters they recognized.

On the initial baseline visit, participants performed a set of RSVP measurements. Each subject was tested with 5 print sizes (defined by the vertical height of a square character configuration, ranging from 1.18 to 1.78 logMAR). Each print size was tested using 5 different exposure durations (i.e. the duration of each sequentially-presented character). Additional exposure durations were added in units of 1.5 times the preceding duration until a spread of recognition accuracies between 20 - 80% were collected. Five trials of each exposure duration were presented for each print size, resulting in a total of at least 125 individual sentences per participant. A psychometric function (cumulative Gaussian distribution) relating exposure duration to character recognition accuracy was fitted. The exposure duration that elicited 55% recognition accuracy for each print size – converted to log characters per minute (log cpm) – was derived. Log print size (minimum angle of resolution, logMAR) and log characters per minute were then fitted to a continuous bilinear function separately for each participant. Reading speed was assumed to increase linearly with increasing print size in this bilinear function until reaching a plateau. The reading speed associated with the plateau determined the participant’s maximum reading speed (MRS), and the smallest print size eliciting the MRS determined the critical print size (CPS)^1,25,27^. The print size corresponding to 0.2 logMAR below the individually fitted CPS and the associated exposure duration were tested in both stimulation sessions^18^, to provide room for any improvement in reading accuracy (see Fig. 2 for an example).

**Fig. 2.**
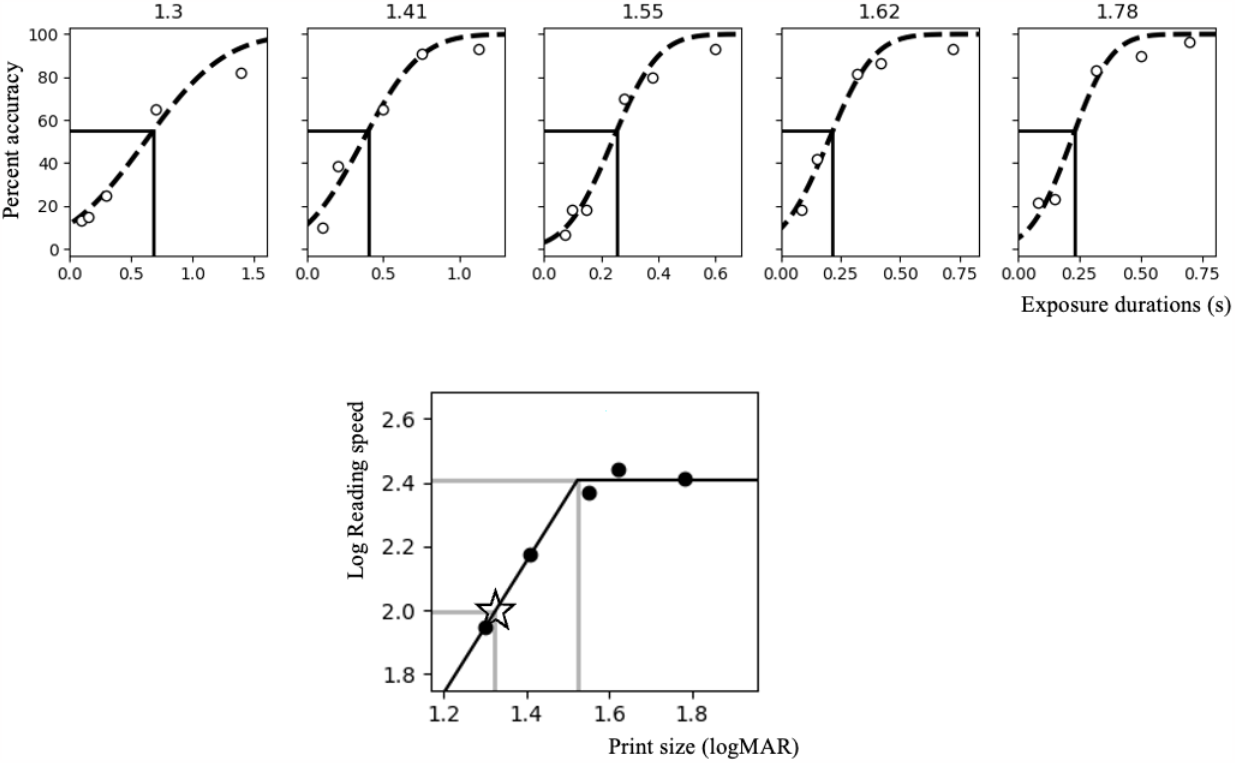
Baseline RSVP curve fitting for a sample subject. The upper panel shows a representative RSVP baseline measurement involving 5 print sizes (1.3, 1.41, 1.55, 1.62, and 1.78 logMAR). The accuracy for each exposure duration is represented by the white circles. The black dotted curve represents the psychometric function (cumulative Gaussian distribution), while the vertical dark line points to the exposure duration that elicits 55% character recognition accuracy. The lower panel shows the bilinear function relating log reading speed (dark circles) to a range of tested print sizes. The upper horizontal and vertical gray lines indicate the maximum reading speed (MRS) and critical print size (CPS) corresponding to 55% accuracy. The lower gray lines indicate the reading speed (in terms of exposure duration) and print size used during the stimulation sessions, which corresponds to a size of 0.2 logMAR smaller than CPS (marked by the star).

### Transcranial direct current stimulation (tDCS)

The tDCS sessions comprised 2 visits separated by at least 48 hours but no more than 7 days. NIBS was delivered using a battery-powered current stimulator (Neuro Device Group S.A. nurostym tES) and two 5cm x 5cm rubber electrodes placed inside saline-soaked sponges. Subjects received either active or sham (placebo) a-tDCS in random order during the 2 visits. Both experimenter and subject were blinded to the stimulation type. The anodal electrode was placed over the primary visual cortex (Oz in the international 10-20 electroencephalogram system), and the cathode was randomly applied over the left or right cheek^18^, secured using a cap. Active a-tDCS involved a direct current ramping up to 2 mA over 30 seconds, continuously delivered for a duration of 20 mins, then ramped down to 0 in 30 seconds. Sham a-tDCS only consisted of the 30-second ramp-up and down periods and no current applied otherwise. During each stimulation session, subjects first read 15 RSVP sentences at the assigned peripheral location with the selected print size and exposure duration before receiving stimulation. Subjects then underwent 20 minutes of stimulation and performed an identical reading test with 15 new sentences. The reading task was again administered 5 mins and 30 mins after the cessation of the stimulation. The outcome measures were the differences in reading accuracy between the baseline and the three post-tests within the same experimental session.

### Statistical analysis

Statistical analysis was performed using GraphPad Prism 9.2.0 for Windows 64-bit, GraphPad Software, San Diego, California USA, www.graphpad.com. The changes in reading accuracy were not significantly different from a normal distribution (Kolmogorov-Smirnov goodness of fit test, p>0.5). Changes in accuracy were compared with a mixed-design analysis of variance (ANOVA) with stimulation type (active vs. sham) and time (during-vs. 5 mins post-vs. 30 mins post-tDCS) as within-subject factors and stimulus location (left vs. inferior) as a between-subject factor. A p-value of less than 0.05 was considered statistically significant.

## Results

Percent differences in reading accuracy were compared among the 3 post-stimulation tests relative to the pre-stimulation performance. No significant effect of stimulation type (F(1, 15)=0.12, p=0.74), time (F(2, 30)=0.43, p=0.65), or stimulus location (F(1, 30)=0.11, p=0.74) was found (Fig. 3). Additionally, the interaction effect among the variables was not statistically significant (F<1.39, p>0.10).

**Fig. 3.**
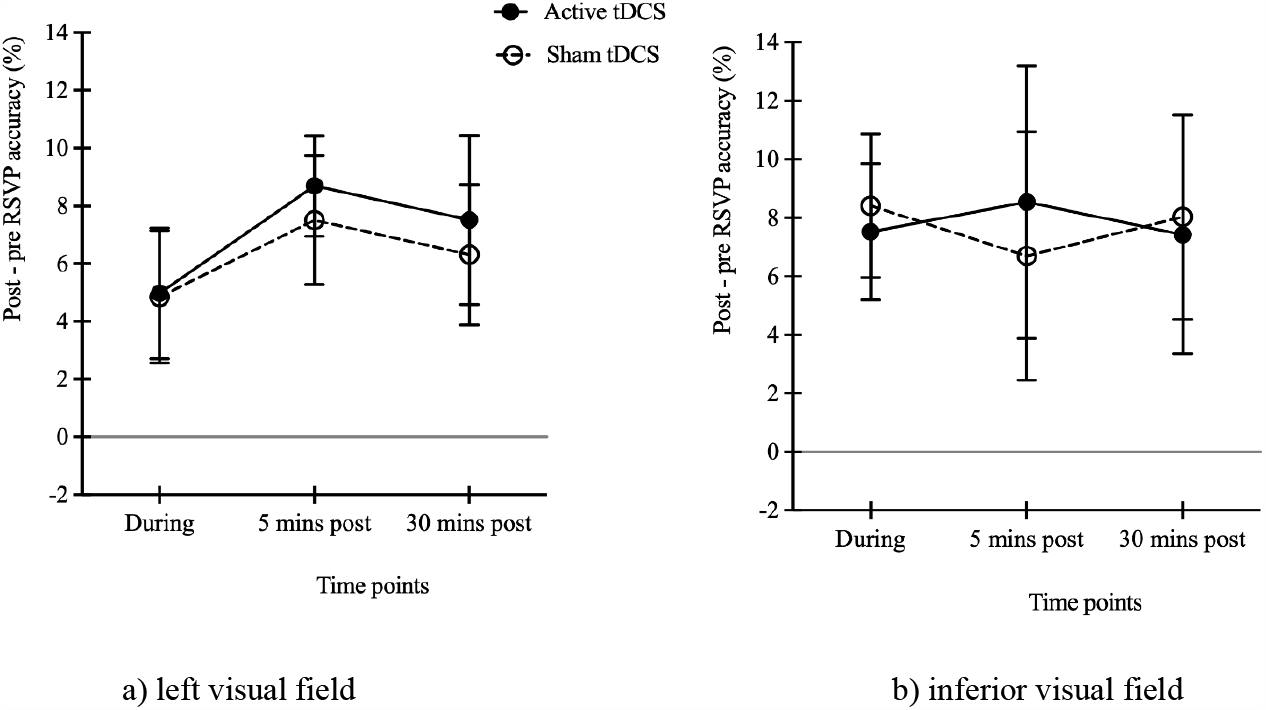
Effect of a-tDCS on RSVP accuracy changes for Chinese characters presented to 10° left and inferior visual fields. Percent differences in RSVP recognition accuracy at each of the 3 post-tests (during-, 5 mins and 30 mins post-stimulation) relative to pre-test at a) 10° left visual field and b) 10° inferior visual field. The black solid and open circles represent the mean differences for active and sham a-tDCS respectively. The error bars are ± 1 standard error of the mean.

## Discussion

The aim of the current study was to investigate the effect of 20 mins of visual cortex active a-tDCS compared with sham a-tDCS on reading individual Chinese characters in a population with normal peripheral vision. In opposition to our hypothesis, we did not find any significant improvement in Chinese reading at peripheral vision following a-tDCS.

Two peripheral locations were examined in the current study as they represent the 2 most frequent para-central regions adopted by AMD patients fixating at a central cross^23^. We did not find a significant difference in accuracy changes when reading Chinese characters presented to these two locations.

Unlike phonetically based languages comprising different numbers of letters with simple spatial forms, Chinese - a logographic orthography language - contains an enormous number of characters (approximately 2500 frequently used Chinese characters^28^) with a wide range of spatial complexities arranged within a square configuration. This unique feature makes Chinese reading in the peripheral vision susceptible to 2 levels of crowding: internal (within-character) and external (between-character)^29^. In the current study, Chinese reading performance was examined using a character-by-character presentation which isolated internal crowding. This is in contrast to studies of English reading in peripheral vision which mainly isolate external crowding between adjacent letters^2,3,18^. Silva et al. found that active a-tDCS increased reading accuracy in patients with macular degeneration who read English sentences presented word-by-word compared with sham stimulation, whereas such improvement was not shown in patients reading Chinese character-by-character^18^. Furthermore, Chen et al. compared the contrast threshold for recognizing either isolated English letters or letters flanked by 2 distractors appearing at 10° eccentricity before and after brain stimulation in normally sighted young subjects. They found that the contrast threshold for recognizing isolated letters was not altered after receiving active a-tDCS. However, a significant improvement was found after the stimulation in the crowded condition^17^. These studies suggest that a-tDCS may have a greater impact on alleviating between-character crowding than within-character crowding, thus enhancing character recognition and peripheral reading performance. Future studies presenting multiple Chinese characters (e.g., recognizing the central character within a trigram) at one time are required to investigate the effect of a-tDCS on internal vs. external crowding and Chinese reading performance in peripheral vision. In addition, only a single session of a-tDCS was adopted in the current study. Exploration of the effects of multiple sessions of a-tDCS on improving Chinese reading performance is warranted.

Recent studies have revealed that another form of NIBS - transcranial random noise stimulation (tRNS) which involves an alternating current that randomly changes in frequency and amplitude, may have a larger effect than a-tDCS on cortical activity^30^ and visual perception^31^. Enhancement of signal-to-noise ratio within the visual cortex due to stochastic resonance has been proposed as the potential mechanism to explain the enhancing effect of tRNS on vision. It is possible that tRNS may improve recognition and reading of Chinese characters presented to peripheral vision. However, this speculation requires further study.

## Conclusions

In agreement with our previous work in a group of patients with central vision loss^18^, active a-tDCS applied to the primary visual cortex did not significantly change Chinese reading performance in normal peripheral vision. While acknowledging the relatively small sample size of the current study which may restrict the generalizability of the results, this exploratory study offers insights into the field of non-invasive brain stimulation and its relationship with reading. Because real-world Chinese reading involves multiple characters presented together, future work should investigate if between-character crowding can be reduced with brain stimulation.

## Acknowledgments

This research was supported in part by grants from The Hong Kong Polytechnic University (internal grant UAG2 to AC), Velux Stiftung Foundation (Grant 1188 to AC and BT) and InnoHK initiative and the Hong Kong Special Administrative Region Government (to AC and BT). Preliminary findings were presented at the Association for Research in Vision and Ophthalmology 2022.

